# BiSulfite Bolt: A BiSulfite Sequencing Analysis Platform

**DOI:** 10.1101/2020.10.06.328559

**Authors:** Colin Farrell, Michael Thompson, Anela Tosevska, Adewale Oyetunde, Matteo Pellegrini

**Affiliations:** Department of Human Genetics, University of California, Los Angeles, CA, USA; Department of Molecular, Cell and Developmental Biology, University of California, Los Angeles, CA, USA

**Author notes:** **Corresponding Author** Matteo Pellegrini.

## Abstract

**Background:** Bisulfite sequencing is commonly employed to measure DNA methylation. Processing bisulfite sequencing data is often challenging due to the computational demands of mapping a low complexity, asymmetrical library and the lack of a unified processing toolset to produce an analysis ready methylation matrix from read alignments. To address these shortcomings, we have developed BiSulfite Bolt (BSBolt); a fast and scalable bisulfite sequencing analysis platform. BSBolt performs a pre-alignment sequencing read assessment step to improve efficiency when handling asymmetrical bisulfite sequencing libraries.

**Findings:** We evaluated BSBolt against simulated and real bisulfite sequencing libraries. We found that BSBolt provides accurate and fast bisulfite sequencing alignments and methylation calls. We also compared BSBolt to several existing bisulfite alignment tools and found BSBolt outperforms Bismark, BSSeeker2, BISCUIT, and BWA-Meth based on alignment accuracy and methylation calling accuracy.

**Conclusion:** BSBolt offers streamlined processing of bisulfite sequencing data through an integrated toolset that offers support for simulation, alignment, methylation calling, and data aggregation. BSBolt is implemented as a python package and command line utility for flexibility when building informatics pipelines. BSBolt is available at https://github.com/NuttyLogic/BSBolt under an MIT license.

## Findings

### Background

DNA methylation, the epigenetic modification of cytosine by the addition of a methyl group to the fifth carbon of the cyclic backbone, is a widely studied epigenetic mark associated with gene regulation[1,2] and numerous biological processes [3–5]. High throughput sequencing combined with bisulfite conversion is a broadly used method for profiling DNA methylation genome wide[6][7]. Treatment of DNA with sodium bisulfite results in unmethylated cytosines being deaminated to uracil, and converted to thymine through PCR amplification, while methylated cytosine, guanine, thymine, and adenine remain unchanged [8]. The methylation status of an individual site or region can be assessed by looking at the number bisulfite converted bases relative to the total number of observed bases. Amongst eukaryotic organisms the majority of genomic cytosines are unmethylated [8–10]. As a consequence, bisulfite sequencing reads originating from the same location but opposite strands are generally no longer complementary. Additionally, when the PCR product of the original bisulfite converted sequence is considered, sequencing reads can be aligned in different orientations within the same strand. Given the asymmetrical nature of bisulfite sequencing libraries and the large number of potential mismatches between the read sequence and the reference the use of a traditional alignment tool would produce low quality alignments.

Bisulfite sequencing alignment tools Bismark[11], BS-Seeker2[11,12], and BWA-Meth[13] successfully adopted a three-base alignment strategy wrapped around established read aligners such as Bowtie2[14,15] and BWA-MEM[14], to accurately align bisulfite sequencing reads. In this strategy, an alignment index or multiple alignment indices are generated against each bisulfite converted reference strand. Relative to the reference, the bisulfite sense strand is the reference with all cytosines converted to thymine and the antisense strand is the reference sequence with all guanines converted to adenine. Before alignment, input reads are *in silico* bisulfite converted so any methylated or incompletely converted bases are converted to remove mismatches relative to the bisulfite reference. Reads are then aligned using the wrapped read alignment tool and the output alignments are integrated together with the original read sequence to form a consensus alignment file. During the generation of a consensus alignment file BS-Seeker2 and Bismark call contextual methylation, where CG methylation is reported distinctly from CH (H=A,C,T) methylation, for every aligned base within an alignment. The regional methylation information provided within alignment calls can provide important context about the epigenetic organization of a genome and the reorganization that occurs in response to disease [16–18]. Methylation calls from aligned reads can also be leveraged to assess the bisulfite conversion status of a read. A high proportion of observed methylated CH sites relative to the total number of observed CH indicates a read that was incompletely bisulfite converted as the majority of CH sites are expected to be unmethylated.

The three base alignment strategy as implemented by BSSeeker2 and Bismark has several limitations. Both tools carry out multiple intermediary alignments to separate alignment indices representing different reference conversion patterns and then integrate intermediate alignments together into a consensus alignment file. Reads with multiple alignments within an intermediate alignment file or across multiple intermediate alignment files are discarded; only reads that align uniquely within a single intermediate alignment are reported. In an effort to reduce the number of reads that align across alignment indices both BSSeeker2 and Bismark have strict default alignment parameters. In addition to being computationally demanding, this implementation can also reduce the number of valid alignments reported, as only the highest quality, unique alignments are output. BWA-Meth resolves this issue by performing alignment to a single bisulfite converted alignment index and processing reads on the fly; but, does not return the read level methylation calls or bisulfite conversion assessment provided by Bismark and BSSeeker2. Additionally, when performing bisulfite sequencing alignment the read conversion pattern is dependent on whether the sequenced DNA fragment is representative of the original DNA sequence or its PCR product. In a directional bisulfite sequencing library only DNA representative of the original DNA fragment is sequenced so the bisulfite conversion pattern is known. In an undirectional library, DNA representative of the original DNA fragment and its PCR product is sequenced so a cytosine to thymine or a guanine to adenine conversion is possible. BS-Seeker2 and Bismark handle undirecitonal libraries by converting input reads using both conversion patterns. This approach doubles the number of reads that must be aligned and generates input reads that will not be represented in the alignment index. BWA-Meth does not support alignment of undirectional libraries.

Here we present BiSulfiteBolt (BSBolt), a bisulfite sequencing platform designed to be fast and scalable while also providing the same read-level methylation calls and quality metrics of BS-Seeker2 and Bismark to preserve compatibility with existing analysis tools. BSBolt alignment is built on a forked version of BWA-MEM[14,19] and HTSLIB[19] with bisulfite specific sequencing logic integrated directly into the alignment process. BSBolt incorporates a pre-alignment read assessment step to assess the correct conversion pattern when aligning undirectional libraries. This eliminates the needs to perform multiple alignments for the same read, improving performance. Additionally, as the output alignment structure is slightly different between each bisulfite alignment wrapper, each tool implements its own methylation calling utility and output format. BSBolt includes a rapid and multi-threaded methylation caller, that outputs methylation calls in CGmap or bedGraph format implemented by BSSeeker2 and Bismark respectively. We show that BSBolt alignments and methylation calling is considerably faster and more accurate than these other bisulfite sequencing alignment wrappers. Additionally, we compare BSBolt to another high performance bisulfite sequencing platform BISCUIT[20]. BISCUIT also incorporates bisulfite specific alignment logic directly into the alignment process, but doesn’t support read level methylation calling or bisulfite conversion assessment during alignment. Despite this, we show that BSBolt offers comparable, or faster, performance. Additionally, to facilitate end to end processing of bisulfite sequencing data BSBolt includes utilities for read simulation utility and aggregation of methylation call files into a consensus matrix.

## Methods

### BSBolt Workflow

#### BSBolt Alignment

BSBolt incorporates bisulfite alignment logic directly within a forked version of BWA-MEM. BSBolt is designed around a single Burrows-Wheeler Transform (BWT) FM-index constructed from both bisulfite converted reference strands. BSBolt utilizes a three base alignment strategy where input reads sequences are fully *in silico* converted before alignment. In this case of undirectional libraries, where a cytosine to thymine or guanine to adenine conversion if possible, BSBolt first analyzes the read base composition. A read, or read pair, with a low proportion of observed cytosines compared to guanine (0.1 by default) will be preferentially aligned with a cytosine to thymine conversion pattern and vice versa. If it is unclear what conversion pattern should be used, both conversion patterns are aligned and the conversion pattern with the highest total alignment score is output. The converted read sequence is aligned using BWA-MEM to the bisulfite FM-index. The resulting alignments are then modified so reads mapping to the sense reference strand are reported as sense reads and the anti-sense reference reported as antisense reads regardless of mapping orientation. The mapping quality of an alignment is assessed by mapping uniqueness using standard BWA-MEM scoring criteria. Additionally, an alignment with alternative alignments on a different bisulfite reference strand is further penalized for being bisulfite ambiguous. Read variation and methylation calls are then made for alignments meeting scoring thresholds using the original read sequence and an unconverted reference sequence. If a difference between the alignment and reference is explainable by bisulfite conversion a methylation call is made for the aligned base; otherwise, reference variation is reported. When calling methylation values, the context of the methylatable base is considered by capturing the local reference context (ie CG or CH). The methylation calls are output as a Sequence Alignment/Map (SAM) flag mirroring the BWA-MEM MD flag. Typically, the majority of CH sites are unmethylated so the expectation is that the majority of CH sites within a read, or read pair, are bisulfite converted. After calling read level methylation this information is leveraged to assess the bisulfite conversion status of the read across all aligned bases within the read, or read pair. The conversion status of the read is conveyed as a SAM flag in the output alignment. Output alignments are then compressed and written to a bam file natively.

#### BSBolt Methylation Calling

BSBolt includes an optimized methylation calling utility that takes advantage of the BSBolt alignment file structure to rapidly call site methylation. The calling procedure proceeds as follows. A read pileup is created using samtools[19], and initialized using pysam[21], for each reference contig with aligned reads. Methylation calls are made for all methylatable bases, or only CG sites, using all reads that pass user specified quality metrics. Methylation values for reference guanine nucleotides are made for reads aligned to the antisense strand and calls for reference cytosine nucleotides are made for reads aligned to the sense strand. This call strategy decreases methylation calling time, as information about the origin strand can be quickly interpreted. Methylation calls are then output in the CGmap file format implemented by BSSeeker2. To aggregate several call files together into a consensus matrix BSBolt includes a rapid and efficient matrix aggregation utility. Bisulfite sequencing techniques often capture methylation sites unevenly, so making a combined matrix of all sites observed across every call file can be inefficient and produce large sparse matrices. BSBolt utilizes an iterative matrix assembly method where individual CGmap files are iterated through to count how often individual sites appear at or above a user specified coverage threshold. If a site is observed in a set proportion of the CGmap files the site is included in the consensus matrix. This process is parallelizable across several threads for efficiency. BSBolt supports output of matrices containing methylation values and counts of methylated and total bases at each site.

#### BSBolt Simulation

BSBolt Simulate utilizes a modified version of WGSIM[22] wrapped with python to simulate bisulfite converted reads with site specific methylation information incorporated across reads. Given a reference sequence global methylation values are set by randomly selecting a methylation value for all methylatable bases depending on context (CG or CH) or by passing a methylation profile in the form of a CGmap file. Reads are then simulated by randomly selecting a genomic position within a reference sequence, sampling the reference sequence at set read length, and insert size for paired end reads, then incorporating sequencing error and genetic variation. The origin strand, and conversion pattern if simulating undirectional reads, is then randomly selected. At every methylatable base within a read the methylation status of the base is set by the probability of observing a methylated base given the reference methylation value. The mapping location, methylation status, and origin bisulfite strand are attached as a fastq comment and output along with the bisulfite converted read sequence and base call qualities. The number of methylated and unmethylated bases covering each methylation site are output as a serialized python object at the end of the simulation.

### Tool Comparisons

BSBolt (v1.4.4), BISCUIT (v0.3.16.20200420), BSSeeker2 (v2.1.8), BWA-Meth (v0.2.2), and Bismark (v0.22.3) were used for comparisons with both real and simulated bisulfite sequencing data. All comparisons were performed on a compute node with XEON X5650 six core (twelve thread) processor (48GB ram) running centos (v6.10). Each tool was provided with 12 compute threads if supported. Default alignment parameters were used unless library specific alignment options were necessary to support the simulated library type. Uncompressed alignment outputs were compressed using samtools (v1.9) before being written to disk. Samtools and BSBolt were provided with two compression threads to minimize any alignment bottlenecks (S. Figure 1). If supported, methylation calls were only made using reads with a mapping quality higher than 20.

### Simulated Bisulfite Library Comparisons

A simulation reference genome was created by sampling approximately 2Mb from each chromosome in the human reference genome (hg38) excluding alternative and sex chromosomes. Briefly, 50bp tiles were randomly sampled from a reference chromosome and included in the simulation reference if the tile contained less than 10 ambiguous bases. The first 10kb of the simulated chr1 was duplicated and added as an additional contig. A series of directional and undirectional bisulfite sequencing libraries were then simulated using BSBolt at various read lengths, read depths, and read qualities with random methylation profiles (Table 1). Alignment and methylation calling tools for each package were compared by aligning a simulation library, sorting the alignment file if necessary, and calling methylation values. Each simulation library was processed by each comparison package sequentially in random order on the same compute node. Read alignments were evaluated by the alignment location and strand. An on-target alignment was defined as a read where 95% of the aligned bases were mapped within the simulated region and mapped to the correct origin strand. An alignment was considered off-target if fewer than 5% of the aligned bases were mapped to the simulation region, the aligned strand of origin was incorrect or flagged as a quality control failure. Accuracy of the CpG methylation calls were evaluated by comparing the called methylation value with the simulated value.

**Table 1:** Simulated Bisulfite Sequencing Library Parameters. The parameters used with BSBolt Simulate to prepare simulation libraries using BSBolt for tool comparisons.

|  |  |  |  |  |
| --- | --- | --- | --- | --- |
| Read Depth | Mutation Rate | Sequencing Error | Sequencing Type | Library Type |
| 20 | 0.005 | 0.005 | Paired End | Directional |
| 20 | 0.005 | 0.005 | Paired End | Directional |
| 20 | 0.005 | 0.005 | Paired End | Directional |
| 30 | 0.005 | 0.005 | Paired End | Undirectional |
| 30 | 0.005 | 0.005 | Paired End | Undirectional |
| 30 | 0.005 | 0.005 | Paired End | Undirectional |
| 20 | 0.005 | 0.005 | Single End | Directional |
| 20 | 0.005 | 0.005 | Single End | Directional |
| 20 | 0.005 | 0.005 | Single End | Directional |
| 30 | 0.005 | 0.005 | Single End | Undirectional |
| 30 | 0.005 | 0.005 | Single End | Undirectional |
| 30 | 0.005 | 0.005 | Single End | Undirectional |
| 8 | 0.005 | 0.005 | Paired End | Directional |
| 8 | 0.005 | 0.005 | Paired End | Directional |
| 8 | 0.005 | 0.005 | Paired End | Directional |
| 8 | 0.005 | 0.005 | Single End | Directional |
| 8 | 0.005 | 0.005 | Single End | Directional |
| 8 | 0.005 | 0.005 | Single End | Directional |
| 8 | 0.01 | 0.02 | Paired End | Directional |
| 8 | 0.01 | 0.02 | Paired End | Directional |
| 8 | 0.01 | 0.02 | Paired End | Directional |

### Targeted Bisulfite Library Comparisons

We next utilized publicly available targeted bisulfite sequencing data (GSE152923) generated from peripheral blood mononuclear cells of four individuals [23].The libraries were generated using the SureSlectXT Methyl-Seq (Aligent) kit and three sequencing libraries were generated for each individual with varying levels of input DNA (1000ng, 300-1000ng, and 150ng-300ng). Each library was sequenced (100bp, paired end) on an Illumina NovaSeq generating an average of 144.1 million (118.5 - 230.5) paired end reads. In addition to the sequencing data, methylation measurements were generated using the Infinium MethylationEPIC array (Illumina) for all four individuals. Whole genome bisulfite alignment indices were generated using hg38 for each bisulfite sequencing package. Every sequencing library was aligned and processed using the same workflow. Alignment files were generated, duplicate reads were marked using samtools (v1.9), and methylation values were called. Each alignment and methylation calling workflow was given a maximum runtime of 288 hours. If an alignment was incomplete at the end of 288 hours, duplicate read marking and methylation calling was performed on the reads aligned during the 288 hour limit. Methylation calls made for CpG sites with more than five reads covering a site were then compared with array methylation values from the same biological sample.

## Results

BSBolt was the fastest alignment tool across all simulation conditions, aligning close to 2.29 million reads per minute on average (Figure 2A). BSBolt was approximately 40% faster than the next fastest alignment tool, BISCUIT. When looking at alignment performance by library type, BISCUIT exhibited similar performance to BSBolt when aligning directional reads, but was approximately 229% slower aligning undirectional libraries (Figure 2A). BSSeeker2, BWA-Meth, and Bismark were slower than both BSBolt and BISCUIT when aligning all library types (Figure 2A). BSBolt and BISCUIT aligned the majority of simulated reads across all conditions (>99%) with high accuracy (>99%). BWA-Meth aligned the majority of reads accurately for directional libraries, but as undirectional libraries are unsupported, BWA-Meth undirectional alignments had low mappability (*μ*=0.724) and a low proportion of aligned reads were on target (*μ*=0.706). BSSeeker2 and Bismark exhibited the lowest average mappability across all simulation conditions at 93.6% and 86.9% respectively but the output alignments were generally accurate (Figure 2B). Moreover, BSSeeker2 and Bismark aligned a low percentage of the simulated reads, 65.3% and 42.4% respectively, when the simulated sequencing error and genetic variation was increased from 0.05% to 2% (S. Table 1). Bismark and BSSeeker2 both discard base call quality information when aligning reads so the low mappability with error prone reads is expected.

**Figure1:**
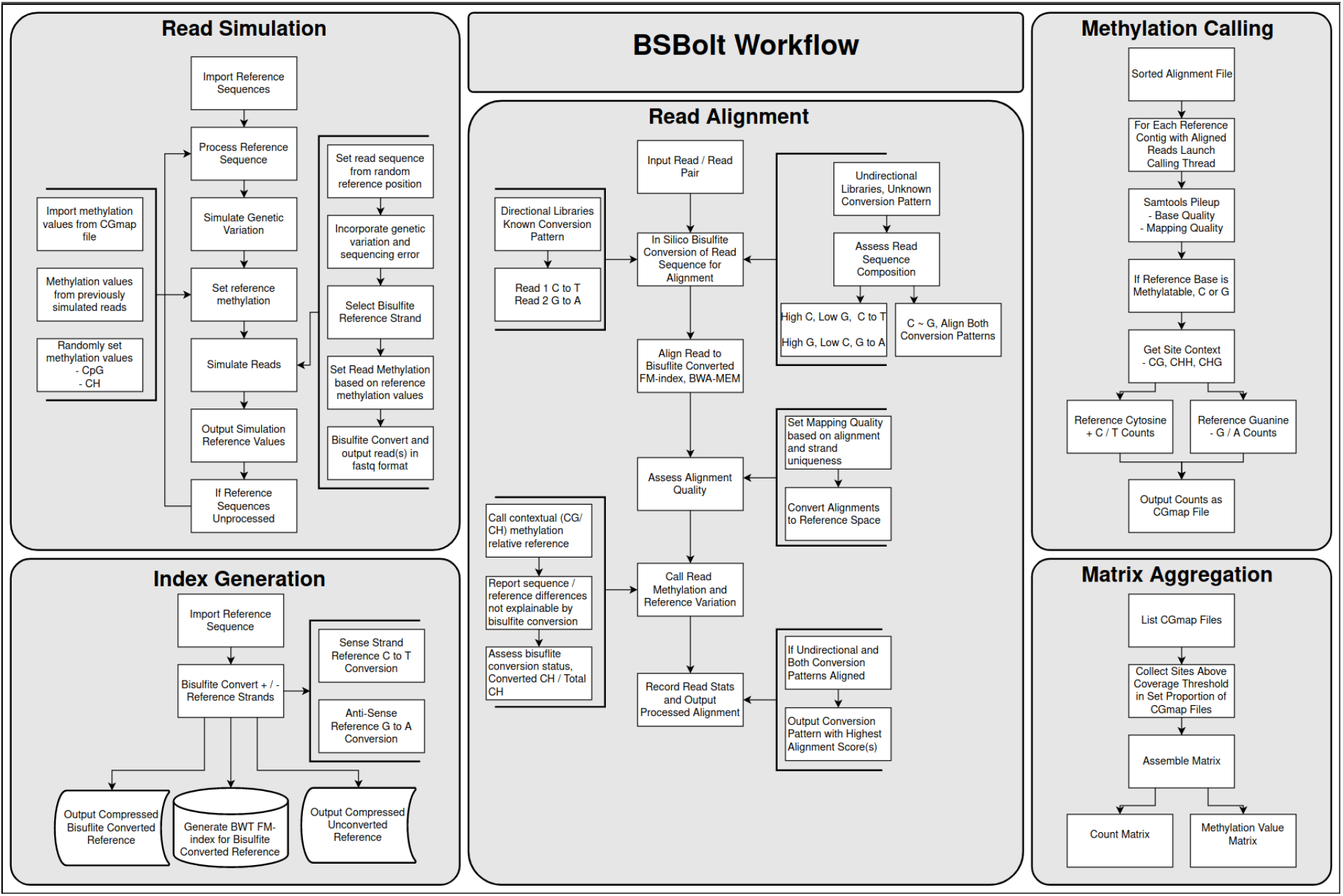
BSBolt Workflows. BSBolt is implemented as a series of discrete modules for read simulation, index generation, read alignment, methylation calling, and matrix aggregation. All BSBolt modules can be run using a command line interface or within a python (>3.6) environment natively.

**Figure 2:**
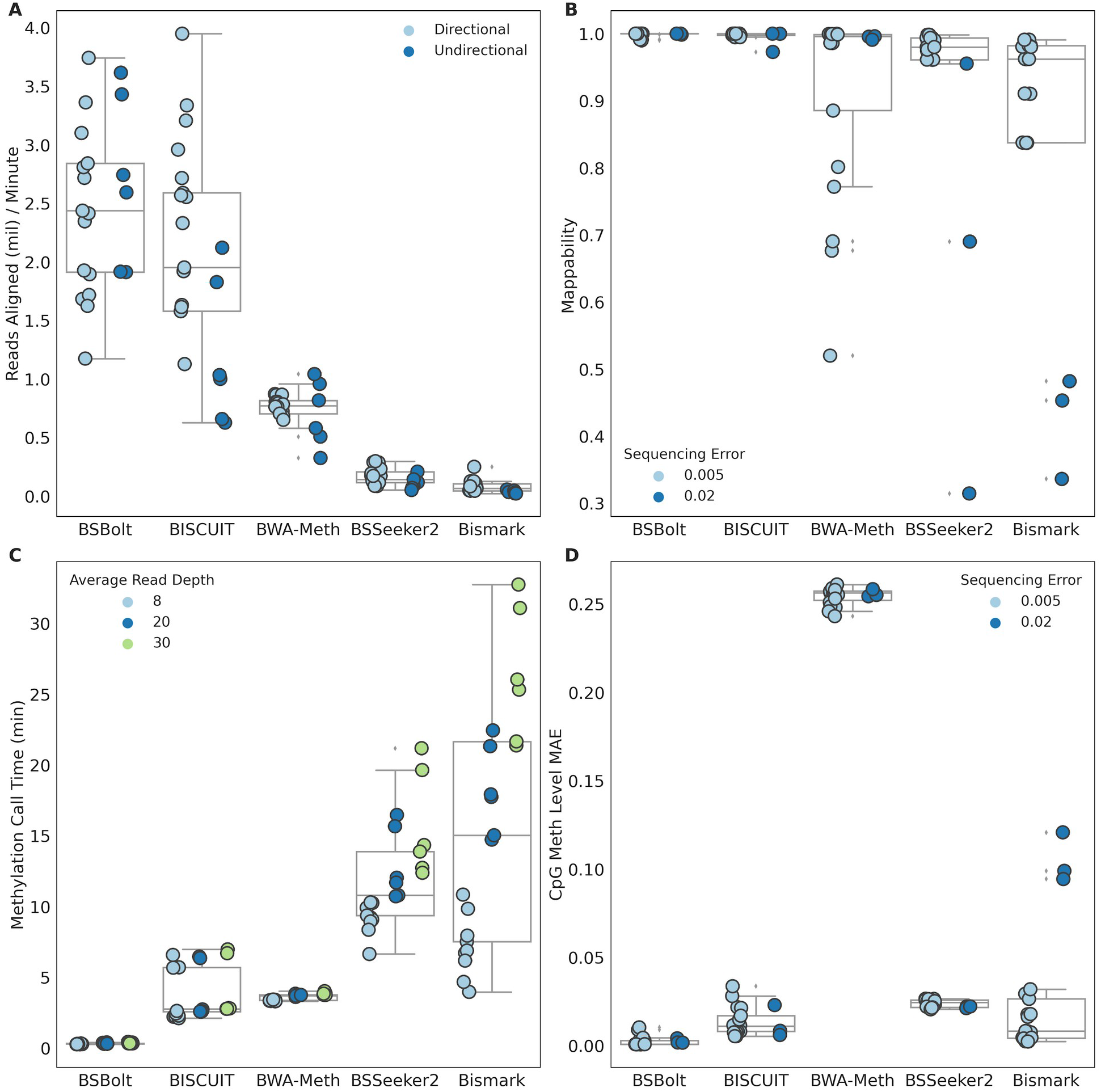
Simulated Bisulfite Sequencing Library Performance. (A) Reads aligned per minute for each bisulfite alignment tool. (B) Proportion of simulated reads mapped during alignments. Note, BWA-Meth does not support undirectional library alignment resulting in low mappability for undirectional libraries. (C) Methylation call time (min) for each alignment tool. (D) Mean Absolute Error (MAE) observed between the simulated and called methylation value.

BSBolt methylation calling was significantly faster than all other tools, with a roughly 11 fold performance advantage over the next fastest tools, BISCUIT and BWA-Meth. BSeeker2 and Bisamark were considerably slower and exhibited a strong relationship between call time and the number of simulated reads (Figure 2C). We also looked at the mean absolute error (MAE) between the number of reads simulated at a given position and the number of reads utilized by each tool to call methylation. BSBolt had the lowest average MAE (0.11 reads) followed by BISCUIT (0.70 reads) and Bismark (0.76 reads). BWA-Meth and BSSeeker2 exhibited high coverage MAE at 6.12 and 8.69 reads respectively. While the BSSeeker2 coverage MAE was high it was not strand biased and the methylation level MAE was small, 0.024. By contrast, the methylation calls made by BWA-Meth were strand biased as shown by the methylation value MAE, 0.255. Overall, BSBolt had the lowest observed methylation level MAE (0.002) followed by BISCUIT (0.013) and Bismark (0.024) (Figure 2D).

The performance of each tool with the targeted bisulfite sequencing libraries largely mirrored the results with the simulation data. However, even though the targeted libraries are directional, BSBolt outperformed BISCUIT aligning an average of 663k reads per minute compared with 637k (Figure 3A). BSSeeker2 failed to align three sequencing libraries within the 288 hour alignment limit, aligning only 78% of reads on average. BSBolt was the fastest methylation calling tool, calling CpG methylation in just 4.35 minutes on average (Figure 3B). We then compared the absolute differences between the sequencing and Illumina EPIC array calls made for the same biological sample, excluding BSSeeker2 alignments as three alignments were incomplete. The absolute differences for all comparisons were combined by tool and binned by effective read coverage, or the number of reads used to call the methylation value (Figure 3C). The called methylation values were highly correlated with the sites called on the EPIC array across all alignment tools (Pearson’s r=.92-98, S. Table 2), as previously reported [24]. Unsurprisingly, as sequencing depth increases the observed mean absolute deviation decreases for all tools. At sequencing depths above 40 reads per CpG BSBolt has the smallest absolute deviation between the sequencing and array calls. Note, due the design of the targeted bisulfite libraries, DNA from one origin strand is preferentially captured over a given region. As a result, the strand bias of the BWA-Meth methylation caller didn’t noticeably impact the methylation calls.

**Figure 3:**
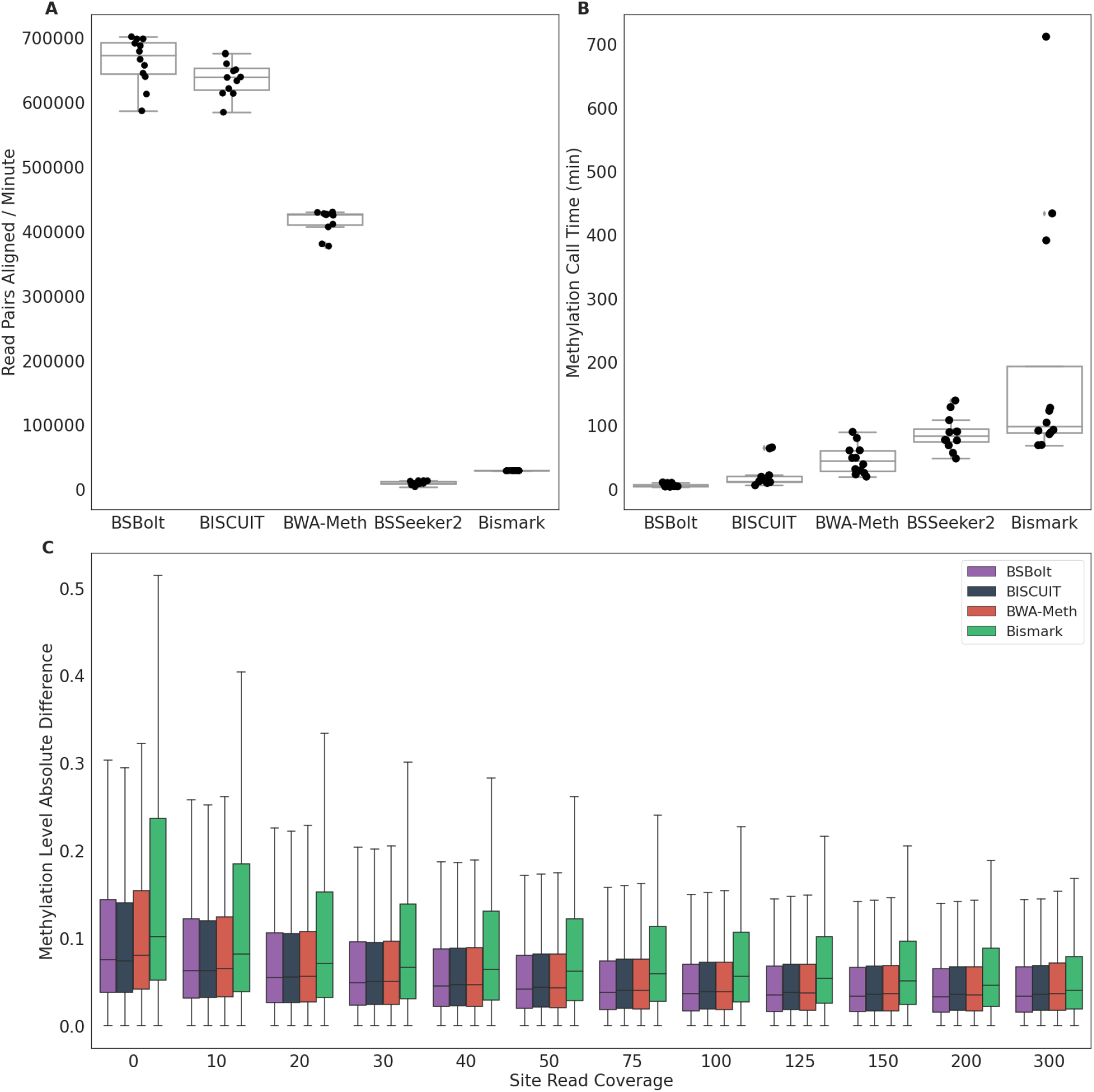
Targeted Bisulfite Sequencing Library Performance. (A) The number of read pairs aligned per minute for each bisulfite alignment tool. (B) Total methylation calling time (min) for each alignment file. (C) The absolute difference between array methylation values and sequencing methylation values for overlapping calls, binned by effective read depth.

## Discussion

Both BSBolt and BISCUIT are significantly faster at bisulfite read alignment while also being more accurate on average than BSSeeker2, Bismark, and BWA-Meth. BSBolt offered marginal performance improvement over BISCUIT with real directional bisulfite libraries, but a large performance gain for the simulated undirectional libraries due to the implementation of a pre-alignment sequencing assessment step. In addition to aligning each read, BSBolt calls contextual read level methylation and assesses read bisulfite conversion, generating alignment information similar to Bismark and BSSeeker2. Importantly, as Bismark and BSSekeer2 have been widely adopted by the community at large it is important to provide the same alignment information to preserve compatibility with downstream tools. BISCUIT offers support for read bisulfite conversion assessment but it is implemented as post-alignment utility.The BSBolt methylation caller was significantly faster than other tools while also providing more accurate methylation calls. Much of this improvement can be attributed to the structuring read alignment before output; by modifying the alignment strand to reflect the bisulfite origin strand methylation calls can be made rapidly without the need to perform additional formatting.

BSBolt is implemented as a python package installable through the python package index[25] and the Anaconda package manager[26].. In addition to a fully command line interface each BSBolt module can be executed natively as an object in a python (>3.6) environment; providing flexibility for informatics pipelines. BSBolt is available at https://pypi.org/project/BSBolt/ and is released under the MIT license.

**Availability and requirements Project name**: BSBolt

**Project home pag**e : https://github.com/NuttyLogic/BSBolt

**Operating system(s)**: Platform Independent

**Programming language**: Python >= 3.6

**Other requirements**: numpy>=1.16.3, tqdm>=4.31.1

**License**: MIT

**RRID**: SCR_019080

## Acknowledgments and Funding

This work was supported by theNational Institutes of Health (T32CA201160 to C.F.).

## Supplementary Information

**Analysis Code**: https://github.com/NuttyLogic/BSBoltManuscript

**Supplemental Table 1**: Simulated Bisulfite Sequencing Library Run Stats

**Supplemental Table 2**: Targeted Bisulfite Alignment Stats

**Supplemental Figure1**: Samtools BAM Conversion Thread Comparisons

**Supplemental Figure2**: BSBolt Performance Characteristics on 150bp Simulated Libraries

## Data Availability

Targeted bisulfite sequencing and EPIC array data deposited in GEO, GSE152923. The pipeline used to simulate bisulfite sequencing libraries is deposited in the analysis repository.

**This work used computational and storage services associated with the Hoffman2 Shared Cluster provided by UCLA Institute for Digital Research and Education’s Research Technology Group.**

---

**Supplemental Figure1:**
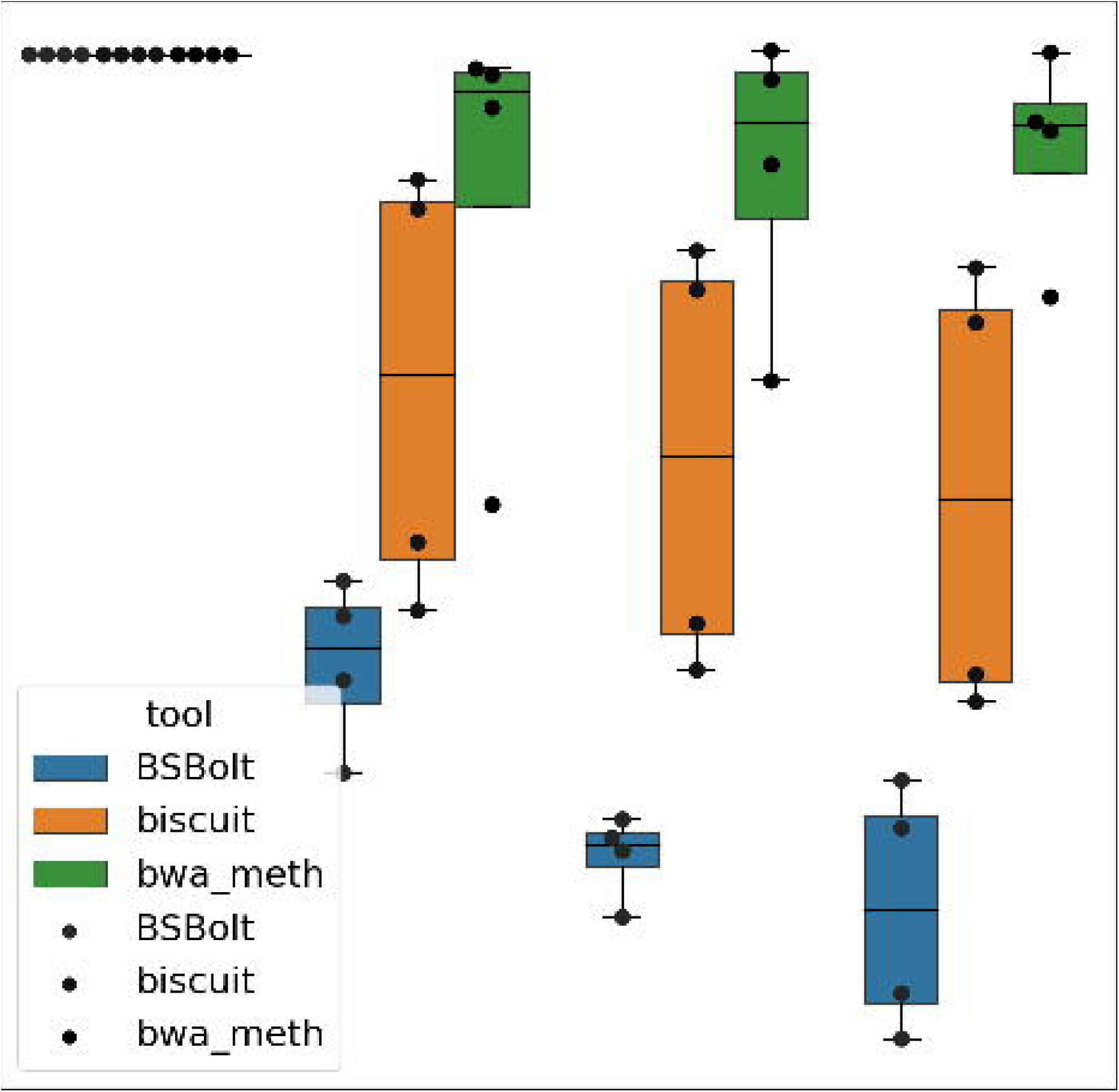
Alignment times for 150 base pair simulated libraries by the number of threads used for SAM to BAM conversion.

**Supplemental Figure 2:**
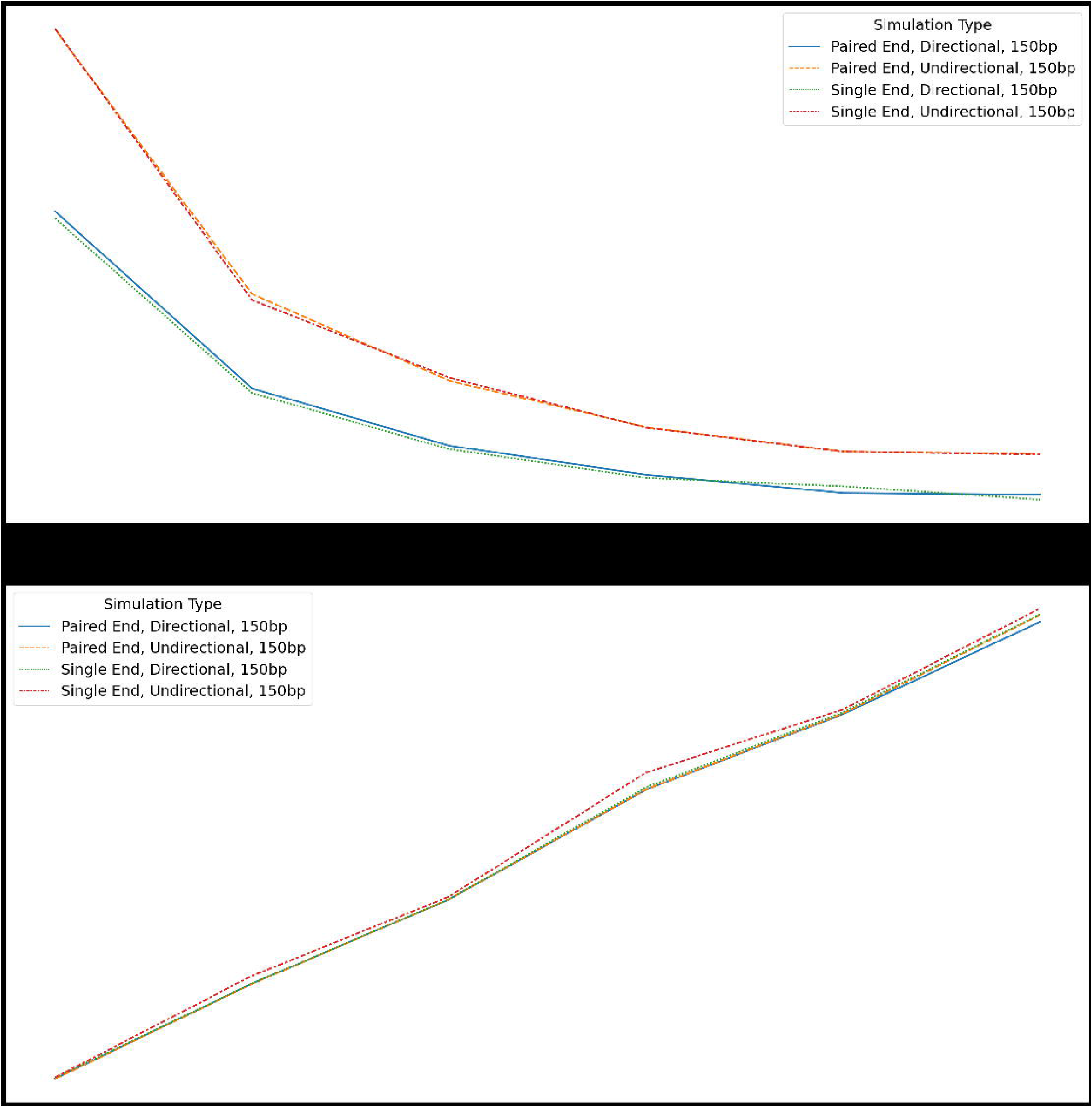
(A) Total alignment time (min) and (B) Maximum memory utilization (mb) for simulated 150 bp bisulfite sequencing libraries by the number of alignment threads provided to BSBolt.

